# Sampling bias and the robustness of ecological metrics for plant–damage-type association networks: Comment

**DOI:** 10.1101/2023.02.14.528448

**Authors:** Sandra R. Schachat

## Abstract

Bipartite network metrics, which link taxa at two trophic levels, are notoriously biased when sampling is incomplete or uneven (Blüthgen et al., 2008; Dormann and Blüthgen, 2017; Fründ et al., 2016). Yet a new contribution (Swain et al., 2023, henceforth SEA) claims the opposite: that bipartite network metrics are minimally sensitive to incomplete sampling and, in fact, perform better at low sample sizes than traditional richness metrics. Here I show that SEA achieved this extraordinary finding by abandoning accepted practices, including practices from the authors’ previous papers.

---

Recently, Currano et al. (2021) compared insect herbivory across fossil angiosperm assemblages. Their study was based on damage type (DT) richness and bipartite network metrics. Currano et al. rarefied data from each fossil assemblage to a uniform level of sampling before conducting their comparisons of damage type richness. They did not report any bipartite network metrics that had been averaged across fossil assemblages, and certainly did not report just one value of each bipartite network metric, averaged across all assemblages under study. Instead, Currano et al. reported metrics separately for each fossil assemblage.

These two practices, of (i) rarefying richness estimates before making comparisons, and (ii) reporting bipartite network metrics separately for each assemblage or site under study, are ubiquitous if not universal.

Currano et al. did not follow these conventions, however, in the Swain et al. (2023) paper they coauthored. SEA examined the sensitivity of the abovementioned measures—richness, and bipartite network metrics—to sampling incompleteness. SEA’s first unprecedented step was to exclusively report unstandardized richness estimates (Figure 1C). Next, SEA evaluated metrics only after normalizing values for each assemblage to equal 1 at the highest sampling threshold, thus removing information about how relative metric values for different assemblages vary with sampling (Figure 1D,E). SEA reported a single value for each bipartite network metric at each level of sampling; their figures illustrate metric values only after they have been averaged across up to 63 assemblages (Figure 1F,G). Had SEA followed their previous practice of rarefying richness estimates, established since insect damage was first compared across fossil localities in the 1990s (Wilf and Labandeira, 1999)^1^, their analyses would have shown richness to be minimally sensitive to sampling incompleteness (Figure 1B). And had SEA reported the value of each bipartite network metric on a per-assemblage basis, as Currano et al. did, it would have been clear that many bipartite network metrics vary considerably with sampling completeness (Figure 1A), as countless ecological studies have previously shown. Therefore, SEA’s conclusion that bipartite network metrics are relatively unbiased by incomplete sampling is an artifact of the decision to upend the authors’ own previous practices for comparing assemblages and reporting their results.

**Figure 1:**
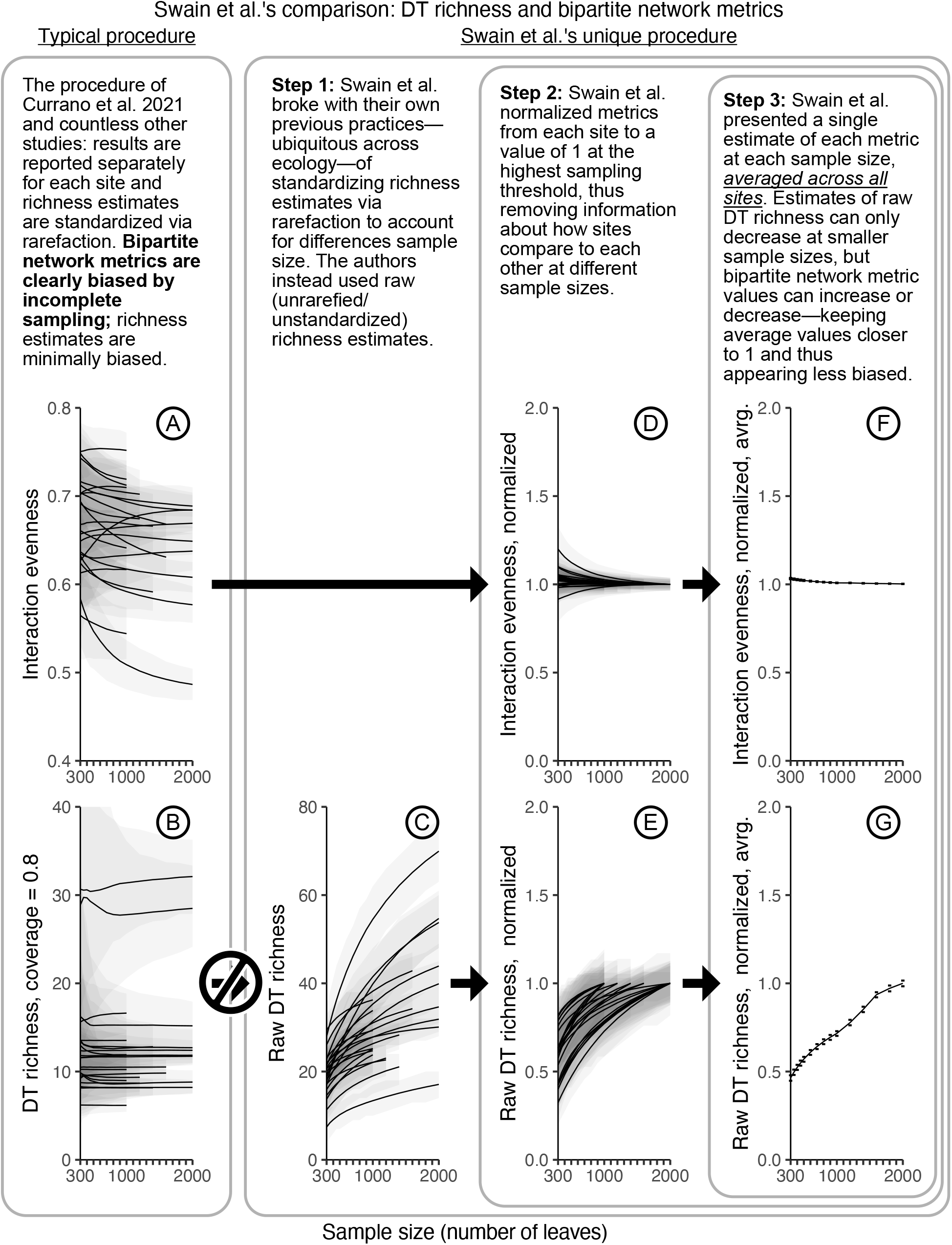
The sensitivity of an exemplar bipartite network metric (interaction evenness) and damage type richness to incomplete sampling, illustrated according to ecological conventions (A, B) and according to SEA’s unique methodology (F, G). The fossil assemblages illustrated here are those from SEA’s compilation with at least 1,000 leaves examined.

## The core issue: Incomplete sampling of fossil herbivory

With either simulated data or data from modern communities, many research teams have found these metrics to be biased by incomplete sampling (e.g., Blüthgen et al., 2006; Rivera-Hutinel et al., 2012; Morris et al., 2014; Fründ et al., 2016; Blüthgen et al., 2008; Dormann et al., 2009). This bias is caused by the inability to differentiate an interaction that genuinely does not occur (a true absence) from an interaction that occurs but was not observed (a failure to detect). A cursory examination of a paleontological dataset illustrates the pitfalls of calculating bipartite network metrics from fossil herbivory data. Such calculations are predicated on the assumption that a dataset is nearly complete, which, in the case of fossil herbivory data, would mean that rarefaction curves for damage type richness on each host plant would approach an asymptote (Figure 2A). In reality, however, many damage types are undetected, even in well-sampled fossil assemblages. Hundreds of angiosperm leaves were examined from the Rose Creek assemblage (Xiao et al., 2022b). Even what is considered good coverage for a fossil dataset is minuscule compared to canonical studies of extant communities. Whereas Xiao et al. (2022a) were only able to examine 244–2,216 cm^2^ of leaf surface area per host plant taxon (Figure 2), Novotny et al. (2012) examined 15,000,000 cm^2^ of leaf surface area per host plant taxon.

**Figure 2:**
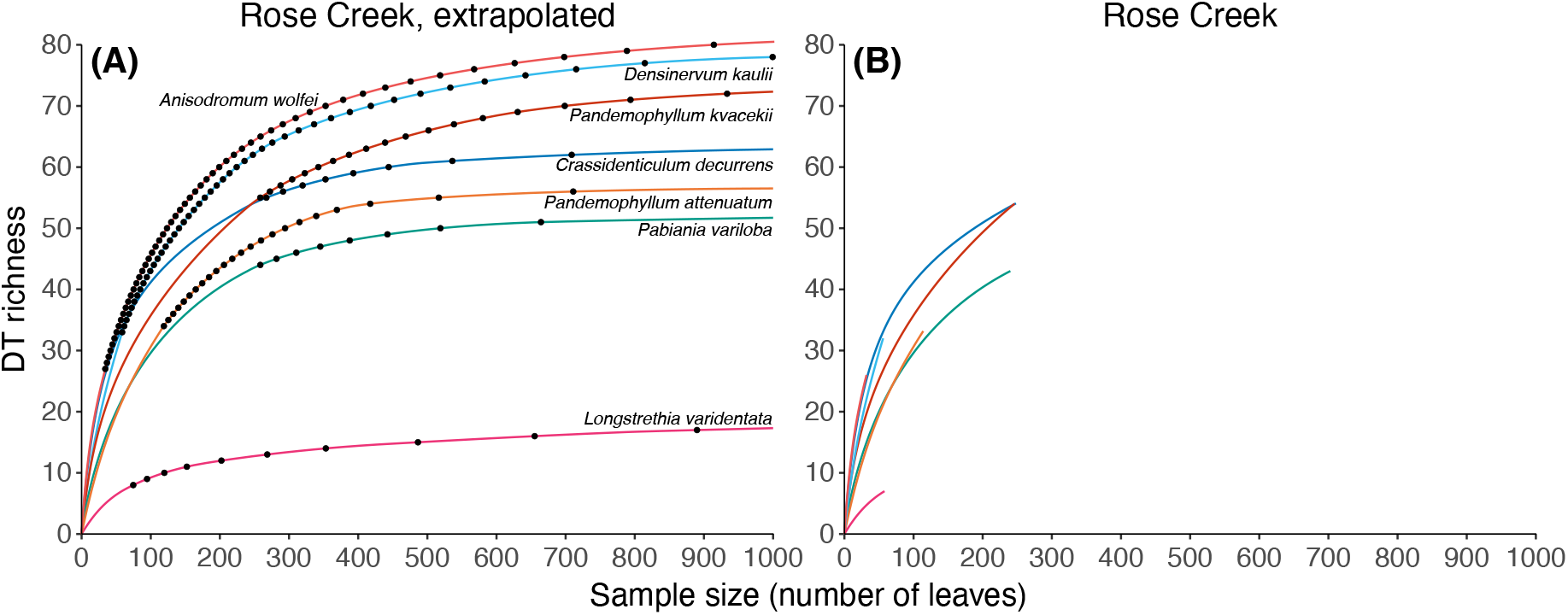
Rarefaction curves for damage type richness on the angiosperm foliar taxa from Rose Creek (Xiao et al., 2022a) represented by 30 or more specimens. The left panel shows extrapolated rarefaction curves in which sampling is nearly complete; black dots indicate interactions that were not detected in the dataset. The right panel shows rarefaction curves from the empirical dataset. N.B.: the data for Rose Creek were published separately from the original paper by Xiao et al. (2022b).

As would be expected with inherently limited fossil sampling, rarefaction curves show that many damage types remain undetected on the seven most abundant host plant taxa at Rose Creek (Figure 2B). In other words, undetected interactions comprise far more than a few rare damage types on a few rare plant taxa. The inability to detect so many interactions will bias estimates of bipartite metrics, because damage types that occur on multiple host plant taxa are likely to be known in fossil datasets from only one such taxon.

## SEA’s unique approach

SEA evaluated sampling bias by examining how sample size impacts a metric’s mean value, averaged across 63 assemblages. In their previous work, however, the same authors reported metrics (bipartite, richness, etc.) for individual assemblages rather than averaged values across dozens of assemblages. Figure 3 shows all bipartite network metrics (other than functional complementarity) illustrated in the SEA’s main text. Here I have illustrated the average value of each metric for each fossil assemblage at each level of sampling; SEA did not illustrate these per-assemblage results in their publication or even in the supplemental material.

**Figure 3:**
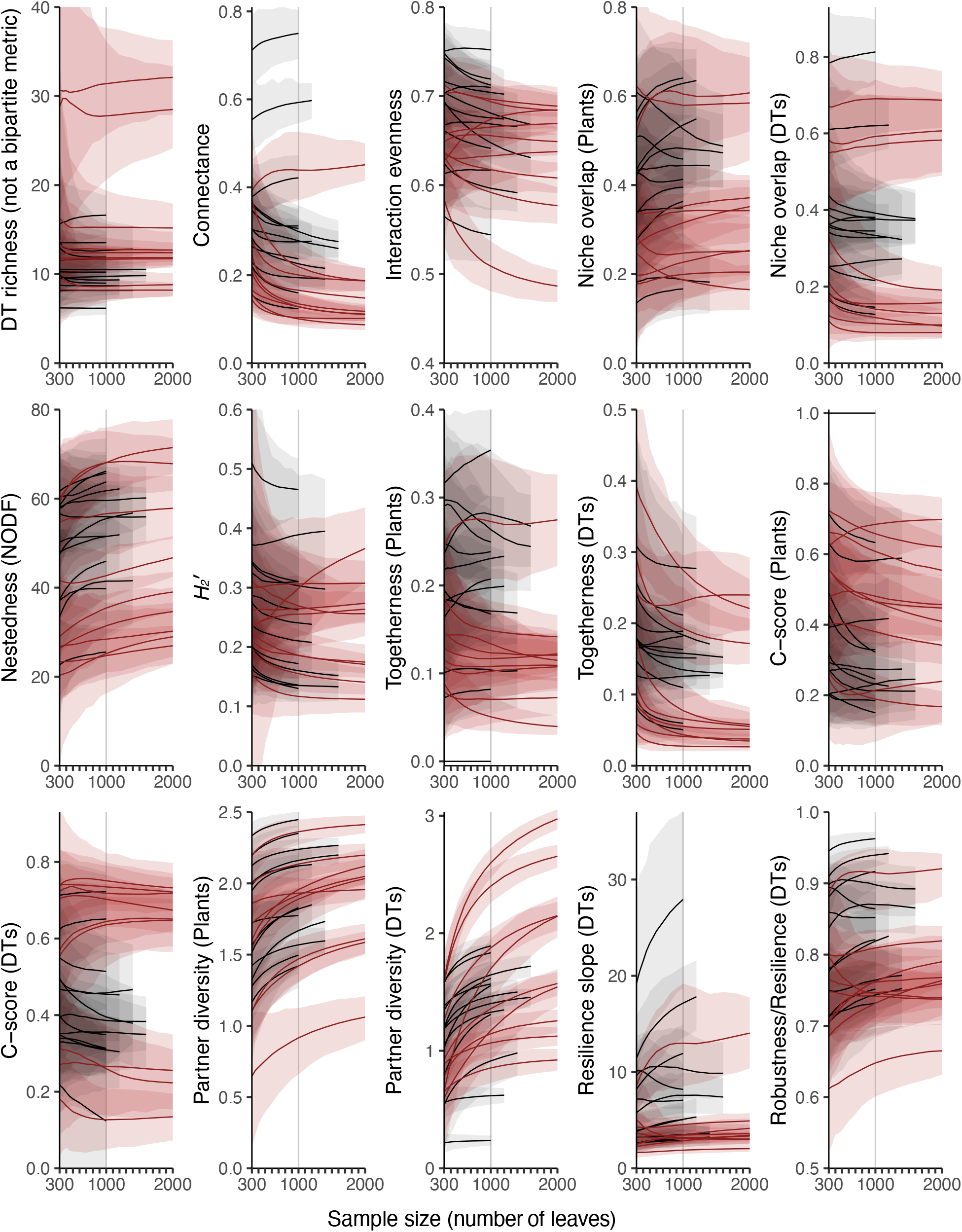
Damage type (DT) richness and the bipartite network metrics reported by SEA, as calculated for 300–2,000 leaves. Each line represents an assemblage in SEA’s data compilation with at least 1,000 leaves examined. Damage type richness was calculated with coverage-based rarefaction. Estimates for each level of sampling represent the average value of 1,000 subsampling iterations at each sampling threshold. More detailed views of all metrics are provided in Figures A1, A2, and A3. Note also that size-based rarefaction, a more traditional method, yields similarly robust results to coverage-based rarefaction (Figure A4). The x-axis begins at 300 leaves (the threshold employed by Currano et al. (2021)), with vertical gray lines at 1,000 leaves (the far more stringent threshold that Currano et al. (2021) suggested all other workers should adhere to).^2^

When the sensitivity of bipartite network metrics is evaluated by visualizing each assemblage separately, it becomes clear that metrics often vary widely with changing sample size (Figure 3). The problem shown in Figure 3, but obscured in SEA’s graphs of average metric values, is the frequency with which the lines representing different fossil assemblages cross each other. If Site A yields a higher value of interaction evenness than Site B when each are represented by 5,000 leaves, but if this pattern is reversed (i.e., Site B yields a higher value for this metric) when the sites are subsampled to 300 leaves each, interaction evenness is not reliable when calculated from only 300 leaves.

## New types of data cannot circumvent bias caused by incomplete sampling

SEA claim that the use of “feeding event occurrence” data will yield improved estimates of bipartite network metrics. Feeding event occurrence data are a red herring in discussions of the bias caused by incomplete sampling, because the fundamental problem is that many interactions remain undetected in fossil datasets. In other words, additional detail about those interactions that have been detected does not compensate for the missing information about the interactions that have not been detected.

When feeding event occurrence data are used in place of traditional data for the Rose Creek assemblage (Xiao et al., 2022a, illustrated here in Figure 2), the quantitative bipartite metrics that SEA illustrated in their Figure 4 appear no less biased by sampling incompleteness (Figure 4).^3^ The number of plant taxa included in the network, rather than the use of feeding event occurrence data, is primary determinant of how biased each metric appears. Because the original dataset being subsampled here is already highly incomplete and uneven (Figure 2), even slight variation in the value of a particular metric suggests that the metric would take on a very different value if calculated from a dataset in which rarefaction curves for each host plant taxon neared an asymptote.

**Figure 4:**
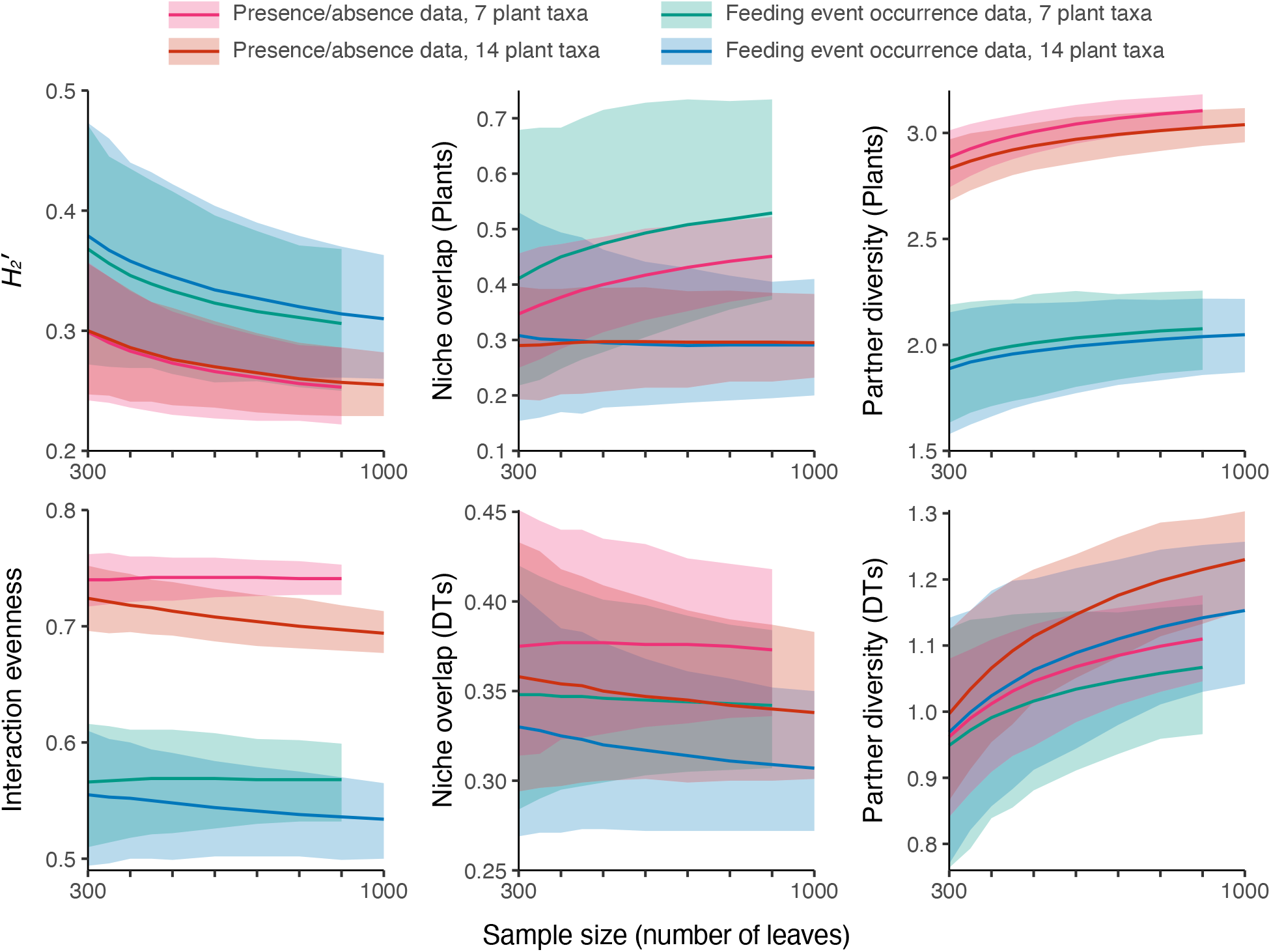
Changes to the values of quantitative bipartite network metrics as sampling becomes more complete, estimated for the Rose Creek assemblage.

Variation in the value of *H* _2_ ′ further underscores this point. *H* _2_ ′ is a bipartite network metric that was devised to be unbiased by incomplete sampling. But, as its authors explained, the uneven sampling that is a hallmark of fossil herbivory datasets remains a major problem such that, “systematic sampling bias (e.g. a taxonomic focus within a guild) will therefore affect the conclusions of comparisons within or across networks” (Blüthgen et al., 2006). In other words, the unevenness of fossil herbivory datasets may be even more difficult to overcome than the incompleteness of these datasets. (SEA did not acknowledge the potential pitfalls of uneven sampling.) The variability of *H*_2_′, of all metrics, when calculated from a relatively large fossil dataset (with or without the use of feeding event occurrence data), indicates that bipartite metrics are not appropriate for fossil data at realistic sample sizes.

## Appendix

### What makes a metric robust?

Because bipartite network analysis is new to many paleobotanists, this discussion begins with two cartoon examples to build intuition. Consider a hypothetical bipartite network metric whose possible values range from 0 to 1. This metric is calculated for five sites. The amount of data collected is denoted by the number of leaves sampled. In Figure A1A, this metric is robust to incomplete sampling, for two reasons. First, the value of this metric for each site changes minimally from the lowest level of sampling (300 leaves) to the highest (5,000 leaves). Second, the relative positions of the five sites do not change with sampling completeness. Depending on the uncertainty surrounding each estimate, at low levels of sampling there may not be enough statistical power to discern significant differences between Sites 2 and 3, and Sites 4 and 5. But there are no false-positive results at low levels of sampling.

Simply put, the lines representing each site have similar slopes that are all nearly horizontal. Which, when sample size is plotted along the x-axis, illustrates robustness to sampling incompleteness.

Figure A1B, however, shows a hypothetical metric that is not robust to sampling incompleteness. The values of this metric for Sites 2–5 change noticeably as sampling becomes more complete. Even more worrying, however, is that the relative values of this metric for the five sites change dramatically as sampling becomes more complete. At the lowest level of sampling, there are clear differences between Sites 2–5. At 300 leaves, the value of this metric is much higher for Site 3 than for Site 2. Similarly, it is much higher for Site 5 than for Site 4. These two findings not only become insignificant, but are in fact reversed, at higher levels of sampling. The differences between Sites 2 and 3, and Sites 4 and 5, are therefore false positives when the analysis is conducted with only 300 leaves.

In other words, the lines in Figure A1B do not have similar slopes. This indicates that the metric in question is not robust to sampling incompleteness.

### Applying these concepts to fossil data

Rarefied estimates of damage type richness are robust to differences in sampling completeness. Coverage-based rarefaction, a method that was established ten years ago (Chao and Jost, 2012) and has been recommended for studies of fossil herbivory (Schachat et al., 2022), yields remarkably consistent richness estimates across sample sizes (Figure A1C). The relative values of the estimates for different assemblages do not change with sample size; *e*.*g*., the Messel assemblage (Wappler et al., 2012) has higher damage-type richness than the Willershausen assemblage (Adroit et al., 2018) regardless of whether the original datasets are subsampled to 5,000 or 300 leaves before being rarefied by sample coverage. The lines in Figure A1C have similar slopes which are nearly horizontal, as is the case for Figure A1A.

In contrast, bipartite network metrics do not fulfill either of the criteria for robustness to incomplete sampling when calculated for the fossil datasets in SEA’s compilation. Estimates for each metric can change considerably as sampling increases, and the relative values for different assemblages vary quite a bit. There are many false positives: noticeable differences among fossil assemblages observed when only 300 leaves are sampled that disappear or are reversed when more leaves are sampled.

Consider the bipartite network metric termed “niche overlap,” calculated for damage types (Figure A1F). This metric, like the other three bipartite network metrics illustrated in Figure A1, ranges from 0 to 1. When only 300 leaves are sampled per assemblage, the Eckfeld assemblage (Wappler et al., 2012) yields a mean estimate of 0.33 (95% CI: 0.16–0.59) and the Messel assemblage, also from the Eocene of Germany, yields a mean estimate of 0.18 (95% CI: 0.12–0.26). The mean estimate for Eckfeld lies well beyond the uncertainty estimate for Messel. However, when sampling increases from 300 to 4,000 leaves, the estimates for the two assemblages are nearly identical: 0.19 (95% CI: 0.15–0.23) for Eckfeld, and 0.17 (95% CI: 0.15–0.19) for Messel. The decrease in the estimate for Eckfeld, and the false-positive result of a clear difference between the two assemblages at 300 leaves, both indicate that niche overlap is not robust to sampling incompleteness. (Of note, Eckfeld and Messel are two of only three fossil assemblages in SEA’s compilation—out of a total of 63—for which at least 4,000 leaves were examined. If more assemblages had been sampled as thoroughly as these, it is entirely possible that more of the differences observed at low sample sizes would turn out to be false positives.)

Similar changes in the values of point estimates, with false-positive differences at low sample sizes, occur when niche overlap is calculated for plants rather than damage types (Figure A1G). When only 300 leaves are sampled per assemblage, the Fifteenmile Creek (Currano et al., 2010) assemblage yields a mean estimate of 0.61 (95% CI: 0.44–0.78) and the Wind River Interior assemblage, also from the Eocene of Wyoming (Currano et al., 2019), yields a mean estimate of 0.42 (95% CI: 0.29–0.55). The mean estimate for Fifteenmile Creek lies beyond the uncertainty estimate for Wind River Interior, and vice versa. However, when sampling increases from 300 to 1,600 leaves, the estimates for the two assemblages are nearly identical: 0.49 (95% CI: 0.39–0.60) for Fifteenmile Creek, and 0.47 (95% CI: 0.39–0.56) for Wind River Interior. The major decrease in the estimate for Fifteenmile Creek, and the false-positive result of a clear difference between the two assemblages at 300 leaves, reinforce that niche overlap is not robust to sampling incompleteness. (In light of the findings discussed above for assemblages with over 4,000 leaves sampled, it is entirely possible that the pattern seen at low sample sizes for Fifteenmile Creek and Wind River Interior would not only disappear, but become reversed, if datasets for these two assemblages reached the sample sizes for Eckfeld and Messel.)

These same two indicators of sensitivity to sampling incompleteness—changing estimates with more sampling, and false-positive results at lower sample sizes—are seen for plenty of other bipartite network metrics when calculated with fossil herbivory data (Figures A1D–E, A2, A3). Whereas estimates of damage type richness are consistent across sample sizes as long as they are calculated with an appropriate method, bipartite network metrics are susceptible to biases caused by incomplete sampling.

**Figure A1:**
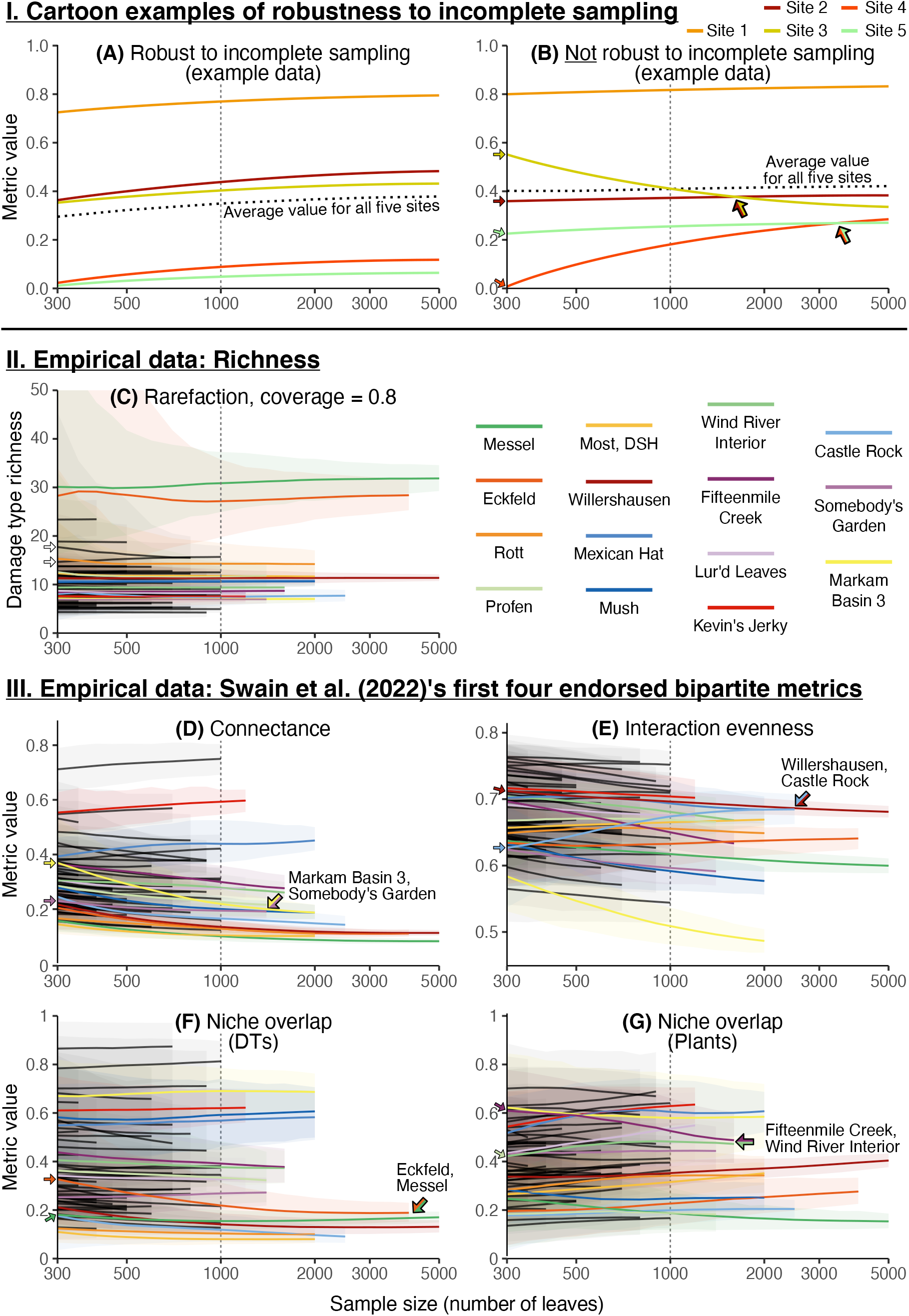
(The sensitivity of various bipartite network metrics to incomplete sampling. For pairs of networks that yield notably different values for a given metric at low levels of sampling, and converge upon similar values at higher levels of sampling, small arrows are placed along the y-axis and larger arrows indicate the points of convergence. C–G use empirical data from the compilation of Currano et al. (2021) and SEA. C shows damage type richness at each assemblage, rarefied with coverage-based rarefaction. D–G show the first four bipartite network metrics mentioned by SEA that were included in their Figures 1–2. The fifteen fossil assemblages that could be subsampled to over 1,000 leaves are color-coded; for the sake of legibility, all other fossil assemblages are in black. A and B show cartoon examples with simulated data. The x-axis ranges from 300 leaves, to which Currano et al. (2021) subsampled datasets for the fossil assemblages in their compilation, to 5,000 leaves, the maximum threshold at which data are available for more than one assemblage. The dotted gray lines represent 1,000 leaves, the minimum level of sampling that Currano et al. (2021) suggest all other researchers adhere to. All x-axes are on a log scale. The cartoon example metrics in A–B, and the bipartite network metrics in D–G, all have possible values ranging from 0 to 1.

#### Bipartite network metrics should not be averaged across 63 fossil assemblages before evaluating sensitivity to sampling incompleteness

SEA presented subsampled data for the same bipartite network metrics shown in Figures A1E–H, A2, and A3. However, rather than presenting estimates for each of the 63 fossil assemblages examined, they presented a single estimate for the value of each metric, at each sampling threshold, averaged across all 63 fossil assemblages. The examples in Figure A1 illustrate why this practice can yield deceptive results. The hypothetical metric illustrated in Figure A1A is minimally sensitive to sampling incompleteness, and, to be sure, the value of this metric averaged across all five sites (shown in the dotted black line) also changes minimally with sampling completeness. However, the hypothetical metric illustrated in Figure A1B also yields an averaged value that varies minimally with sampling completeness, despite the values of this metric for each individual site being highly sensitive to incomplete sampling. When the colored lines in Figure A1B cross as sampling becomes more complete, this indicates a false positive result caused by insufficient sampling. However, because the value of this metric decreases for Site 3 and increases for Site 4 as sampling becomes more complete, the averaged value for these two metrics remains nearly constant despite the values of this metric changing dramatically for the individual sites.

Averaged metric values across multiple sites or fossil assemblages are of minimal relevance to ecological practice. Currano et al. (2021), the only other study that calculated bipartite network metrics for the datasets presented by SEA, reported those metrics for individual networks rather than averaged values. The same is true for neontological studies that reported bipartite network metrics. Averaged or median values of bipartite metrics across all networks under consideration are rarely mentioned, and are reported in addition to—rather than in place of—the values for individual networks (Kaartinen et al., 2010; Schleuning et al., 2016).

Along the same lines, previous studies that examined the sensitivity of bipartite network metrics to sampling incompleteness also reported separate results for each site under consideration (Blüthgen et al., 2006; Rivera-Hutinel et al., 2012; Morris et al., 2014; Fründ et al., 2016). Speaking of which, those previous studies that examined the same questions as SEA used much more complete datasets, and reached the opposite conclusion: that these metrics are rather sensitive to sampling incompleteness. Blüthgen et al. (2008) wrote, “both unweighted network metrics (connectance, nestedness, and degree distribution) and weighted network metrics (interaction evenness, interaction strength asymmetry) are strongly constrained and biased by the number of observations.” Dormann et al. (2009) wrote, “most indices [bipartite network metrics] are strongly affected by network dimensions and sampling intensity” Rivera-Hutinel et al. (2012) wrote, “Except for the robustness to plant loss, all of the network metrics examined here were sensitive to some extent to sampling effort.” Morris et al. (2014) wrote, “the strong association between network size and structure implies that many apparent spatial and temporal variations in network structure may prove to be artefacts.” Fründ et al. (2016) wrote, “we confirmed the high potential for specialization bias.” Dormann and Blüthgen (2017) wrote, “Both incomplete sampling and focal group bias have emerged as methodological concerns.”

Moreover, a study that was designed to overcome the limitations of empirical datasets when evaluating bipartite network metrics’ sensitivity to sampling completeness (Fründ et al., 2016) found that six of SEA’s ten endorsed bipartite network metrics (those featured in SEA’s Figure 4, other than functional complementarity: connectance, niche overlap, C score, nestedness [NODF], resilience slope/extinction slope, robustness) did not meet either of the two criteria for metrics that are robust to sampling incompleteness. The findings of Fründ et al. (2016) provide reason to suspect that SEA’s endorsement of these six bipartite network metrics is an artifact of the incomplete datasets used for their analyses, rather than any inherent properties of these metrics.

#### Additional unresolved issues in applying bipartite network analysis to fossil herbivory data

The first contribution that addressed the appropriateness of bipartite network analysis for fossil herbivory data, Schachat et al. (2023), raised additional issues that were not fully addressed by SEA. The first of these issues is “metric hacking” *sensu* Webber et al. (2020): the opportunistic selection of a subset of the dozens of available bipartite network metrics. The Currano et al. (2021) publication, which is the only contribution before that of SEA to have calculated bipartite network metrics for the assemblages evaluated here, presented generalized linear models involving a number of “selected network properties.” A bipartite network metric called “vulnerability” was included in those models but was not evaluated by SEA; bipartite network metrics called “togetherness,” “resilience slope,” and “robustness” were evaluated by SEA but were not included in the previous generalized linear models (Currano et al., 2021, Table S4).

Furthermore, as Fründ et al. (2016) pointed out, “Many network descriptors relate directly or indirectly to specialization.” But, as detailed by Schachat et al. (2023), the concepts of generalization and specialization become muddied when techniques that were intended for use with taxonomic data (such as bipartite networks, which can link host plant taxa to herbivorous insect taxa) are repurposed for use with functional data. Damage types are functional data: a single damage type observed on multiple specimens can be induced by a variety of distantly-related insect taxa—most if not all of whom may be specialists. This is precisely what Carvalho et al. (2014) found, with specialist insects belonging to various orders convergently inducing Damage Type 12 at two different sites. It appears that SEA were attempting to justify their use of functional data with methods that were intended for taxonomic data when they wrote, “A shift from taxonomic, or species-specific studies, to functional studies, especially studies involving functional traits, has shown the importance of ecological drivers on interactions in both the modern and paleontological ecological literature (e.g., Kunstler et al. 2016; Perez et al. 2020; Barnosky et al. 2017; Schellenberger Costa et al. 2018).” Although the importance of functional data in general, and damage type data in particular, is not in doubt, it is worth noting that none of the four contributions cited by SEA used functional data as inputs for the calculation of network metrics, nor did any of those contributions advocate for such an approach.

## Methods

The datasets analyzed by SEA were downloaded from github. Each dataset was subsampled to each sampling threshold employed by SEA 1,000 times. The following were calculated for each subsampled dataset: the bipartite network metrics reported by SEA, rarefied damage type richness at 300 leaves, and rarefied damage type richness at sample coverage of 0.8.

Swain et al. (2022) reported that host plants represented by fewer than five specimens were not included in calculations of bipartite metrics. Therefore, before subsampling each dataset, any host plant taxa represented by fewer than five specimens were removed. After each iteration of the subsampling procedure, any host plant taxa that were represented by fewer than five specimens in the subsampled dataset were removed.

**Figure A2:**
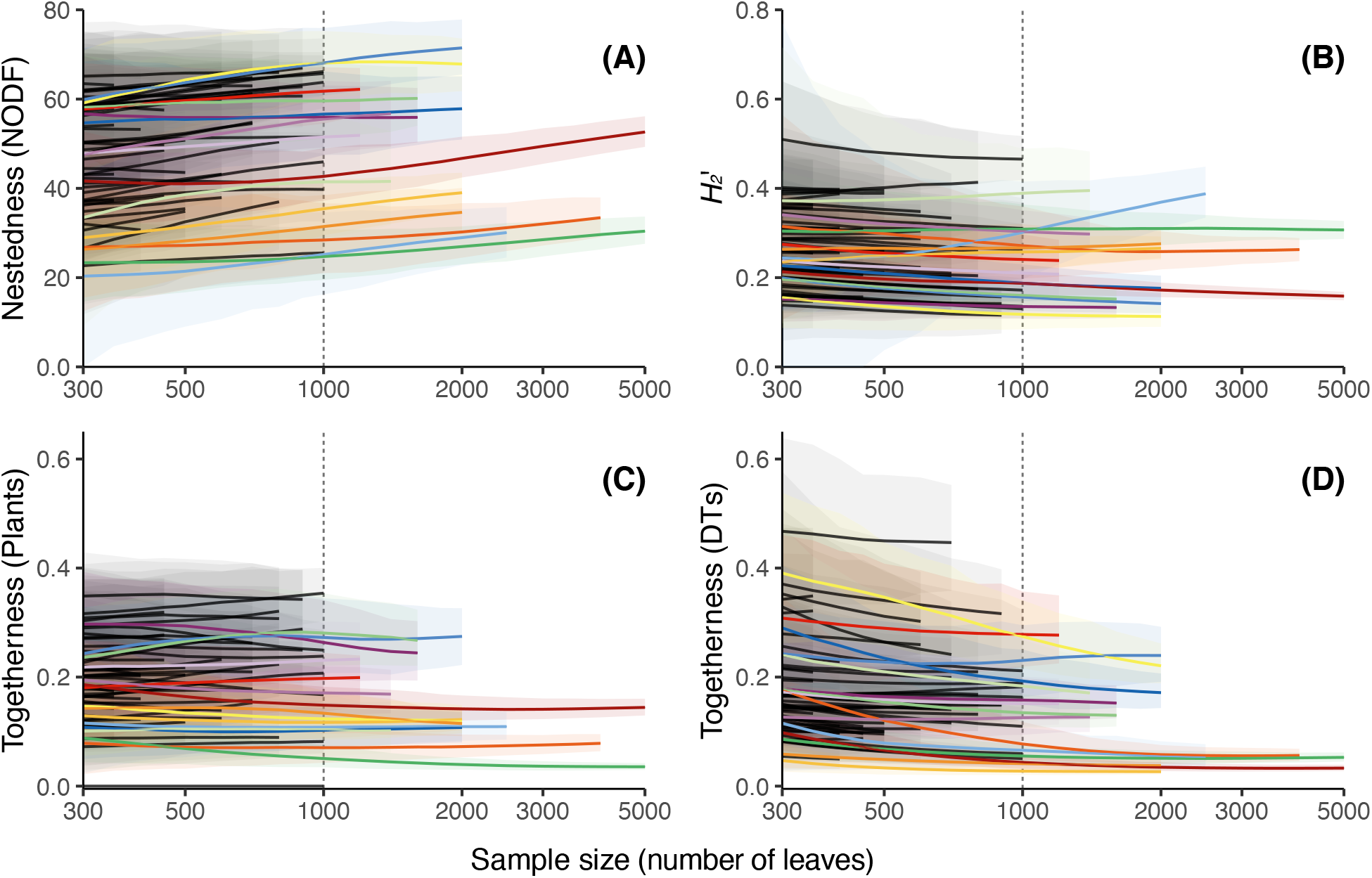
The bipartite networks presented in Figures 1–2 of SEA, continued from Figure A1. For a legend and more information on the scaling of the x-axis, see Figure A1.

**Figure A3:**
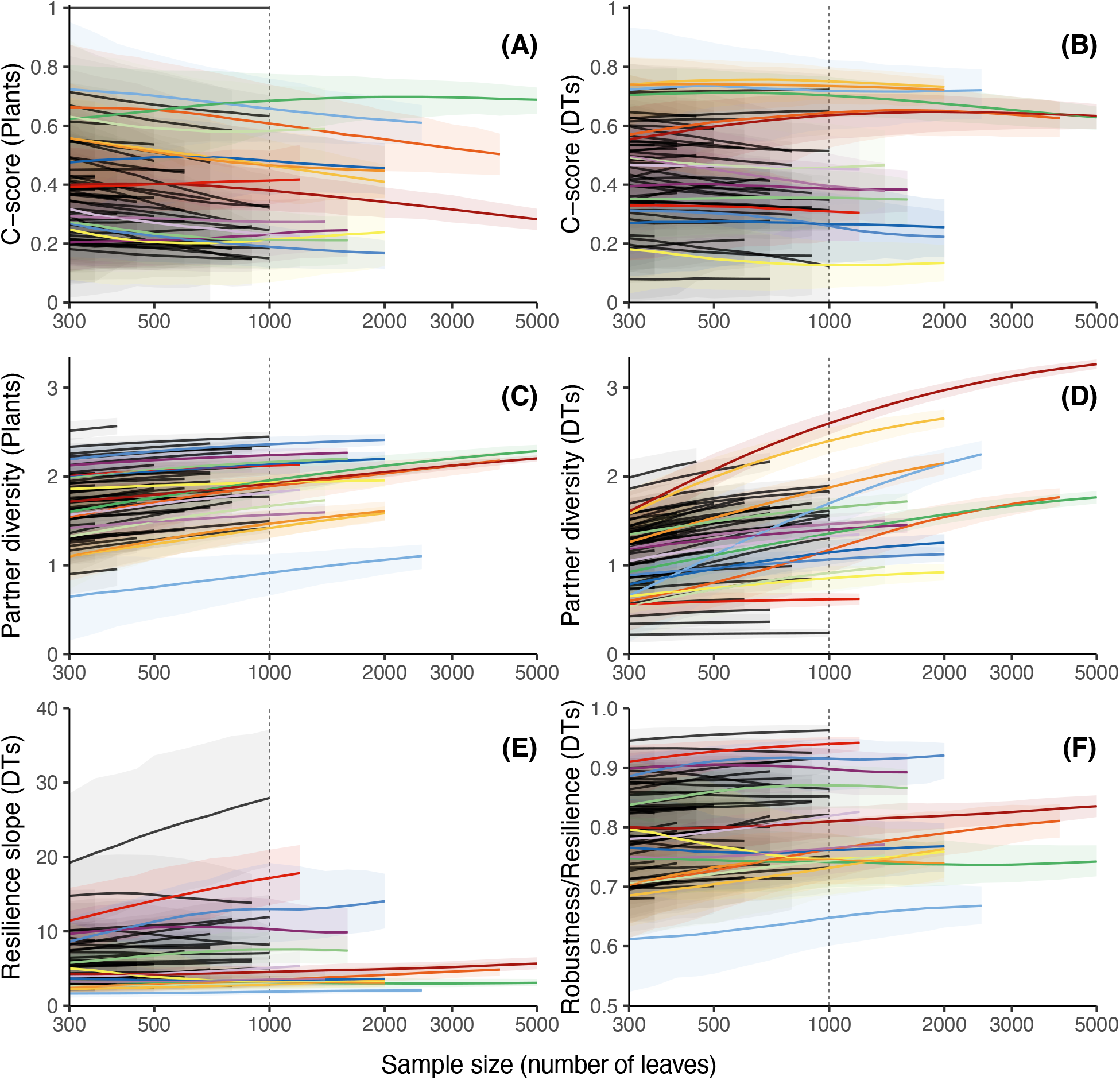
The bipartite networks presented in Figures 2–3 of SEA, continued from Figure A2. For a legend and more information on the scaling of the x-axis, see Figure A1.

**Figure A4:**
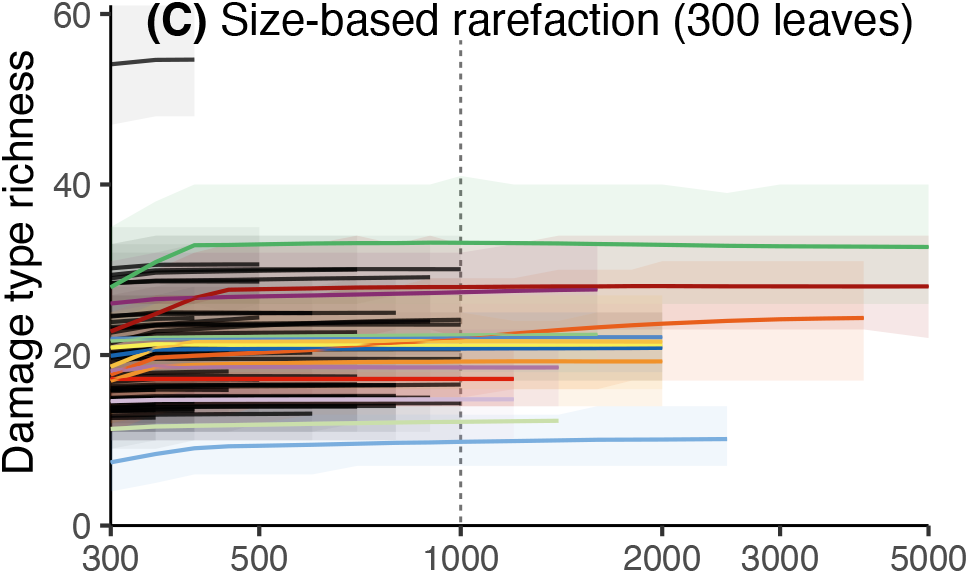
Damage type richness at each fossil assemblage in the compilation of Currano et al. (2021) and SEA, rarefied to 300 leaves, plotted according to the intermediate sample size to which each assemblage was initially downsampled. For a legend and more information on the scaling of the x-axis, see Figure A1.

## Footnotes

1 SEA, too, rarefied their data before reporting richness estimates in their own previous work. See: Azevedo Schmidt et al. (2019) Figs. 5, 6; Currano et al. (2008) Table 1, Figs. 2, 3; Currano (2009) Tables 1, 2, 3, Figs. 4, 5, 6; Currano et al. (2010) Figs. 5, 6, 7; Currano et al. (2011) Figs. 4, 6; Currano et al. (2019) Table 2, Fig. 6; Currano and Jacobs (2021) Table 2, Figs. 4B, 5C,D, 6, 7B; Maccracken and Labandeira (2020) Fig. 6; Maccracken et al. (2022) Fig. 16.

2 “We included in our compilation all sites that were collected in an unbiased manner (i.e., all identifiable leaves scored for herbivory) and had at least 300 dicot leaves. We chose this value to maximize the number of sites that could be included in the meta-analysis and thus spatial and temporal coverage, but we reiterate that new work should target 1000 specimens per census.” Currano et al. (2021), page 4.

3 SEA acknowledge that functional complementarity is, from one point of view, highly biased by incomplete sampling. They say nothing of the sort about any of the other metrics featured in their Figure 4. Of those metrics, the majority (connectance, NODF, togetherness, C-score, robustness/resilience, resilience slope/extinction slope) are “binary” rather than “quantitative”— meaning that each interaction is coded as simply present or absent when the metrics are calculated, ignoring the data on the frequency of each interaction. Binary metrics, therefore, will yield the same result whether or not they are calculated with feeding event occurrence data.

## References

Adroit, B., Girard, V., Kunzmann, L., Terral, J.-F., and Wappler, T. (2018). Plant–insect interactions patterns in three European paleoforests of the late-Neogene—early-Quaternary. PeerJ, 6:e5075.

Azevedo Schmidt, L. E., Dunn, R. E., Mercer, J., Dechesne, M., and Currano, E. D. (2019). Plant and insect herbivore community variation across the Paleocene–Eocene boundary in the Hanna Basin, southeastern Wyoming. PeerJ, 7:e7798.

Barnosky, A. D., Hadly, E. A., Gonzalez, P., Head, J., Polly, P. D., Lawing, A. M., Eronen, J. T., Ackerly, D. D., Alex, K., Biber, E., Blois, J., Brashares, J., Ceballos, G., Davis, E., Dietl, G. P., Dirzo, R., Doremus, H., Fortelius, M., Greene, H. W., Hellmann, J., Hickler, T., Jackson, S. T., Kemp, M., Koch, P. L., Kremen, C., Lindsey, E. L., Looy, C., Marshall, C. R., Mendenhall, C., Mulch, A., Mychajliw, A. M., Nowak, C., Ramakrishnan, U., Schnitzler, J., Das Shrestha, K., Solari, K., Stegner, L., Stegner, M. A., Stenseth, N. C., Wake, M. H., and Zhang, Z. (2017). Merging paleobiology with conservation biology to guide the future of terrestrial ecosystems. Science, 355(6325):eaah4787.

Blüthgen, N., Fründ, J., Vázquez, D. P., and Menzel, F. (2008). What do interaction network metrics tell us about specialization and biological traits. Ecology, 89(12):3387–3399.

Blüthgen, N., Menzel, F., and Blüthgen, N. (2006). Measuring specialization in species interaction networks. BMC Ecology, 6(1):9.

Carvalho, M. R., Wilf, P., Barrios, H., Windsor, D. M., Currano, E. D., Labandeira, C. C., and Jaramillo, C. A. (2014). Insect leaf-chewing damage tracks herbivore richness in modern and ancient forests. PLOS ONE, 9(5):e94950.

Chao, A. and Jost, L. (2012). Coverage-based rarefaction and extrapolation: standardizing samples by completeness rather than size. Ecology, 93(12):2533–2547.

Currano, E. D. (2009). Patchiness and long-term change in early Eocene insect feeding damage. Paleobiology, 35(4):484–498.

Currano, E. D., Azevedo-Schmidt, L. E., Maccracken, S. A., and Swain, A. (2021). Scars on fossil leaves: An exploration of ecological patterns in plant–insect herbivore associations during the Age of Angiosperms. Palaeogeography, Palaeoclimatology, Palaeoecology, 582:110636.

Currano, E. D. and Jacobs, B. F. (2021). Bug-bitten leaves from the early Miocene of Ethiopia elucidate the impacts of plant nutrient concentrations and climate on insect herbivore communities. Global and Planetary Change, 207:103655.

Currano, E. D., Jacobs, B. F., Pan, A. D., and Tabor, N. J. (2011). Inferring ecological disturbance in the fossil record: A case study from the late Oligocene of Ethiopia. Palaeogeography, Palaeoclimatology, Palaeoecology, 309(3-4):242–252.

Currano, E. D., Labandeira, C. C., and Wilf, P. (2010). Fossil insect folivory tracks paleotemperature for six million years. Ecological Monographs, 80(4):547–567.

Currano, E. D., Pinheiro, E. R. S., Buchwaldt, R., Clyde, W. C., and Miller, I. M. (2019). Endemism in Wyoming plant and insect herbivore communities during the early Eocene hothouse. Paleobiology, 45(3):421–439.

Currano, E. D., Wilf, P., Wing, S. L., Labandeira, C. C., Lovelock, E. C., and Royer, D. L. (2008). Sharply increased insect herbivory during the Paleocene–Eocene Thermal Maximum. Proceedings of the National Academy of Sciences of the United States of America, 105(6):1960–1964.

Dormann, C. F. and Blüthgen, N. (2017). Food webs versus interaction networks: Principles, pitfalls, and perspectives. In Moore, J. C., de Ruiter, P. C., McCann, K. S., and Wolters, V., editors, Adaptive Food Webs, pages 9–18. Cambridge University Press.

Dormann, C. F., Fründ, J., Blüthgen, N., and Gruber, B. (2009). Indices, graphs and null models: analyzing bipartite ecological networks. The Open Ecology Journal, 2(1):7–24.

Fründ, J., McCann, K. S., and Williams, N. M. (2016). Sampling bias is a challenge for quantifying specialization and network structure: lessons from a quantitative niche model. Oikos, 125(4):502–513.

Kaartinen, R., Stone, G. N., Hearn, J., Lohse, K., and Roslin, T. (2010). Revealing secret liaisons: DNA barcoding changes our understanding of food webs. Ecological Entomology, 35(5):623–638.

Kunstler, G., Falster, D., Coomes, D. A., Hui, F., Kooyman, R. M., Laughlin, D. C., Poorter, L., Vanderwel, M., Vieilledent, G., Wright, S. J., Aiba, M., Baraloto, C., Caspersen, J., Cornelissen, J. H. C., Gourlet-Fleury, S., Hanewinkel, M., Herault, B., Kattge, J., Kurokawa, H., Onoda, Y., Peñuelas, J., Poorter, H., Uriarte, M., Richardson, S., Ruiz-Benito, P., Sun, I.-F., Ståhl, G., Swenson, N. G., Thompson, J., Westerlund, B., Wirth, C., Zavala, M. A., Zeng, H., Zimmerman, J. K., Zimmermann, N. E., and Westoby, M. (2016). Plant functional traits have globally consistent effects on competition. Nature, 529(7585):204–207.

Maccracken, S. A. and Labandeira, C. C. (2020). The Middle Permian South Ash Pasture assemblage of north-central Texas: Coniferophyte and gigantopterid herbivory and longer-term herbivory trends. International Journal of Plant Sciences, 181(3):342–362.

Maccracken, S. A., Miller, I. M., Johnson, K. R., Sertich, J. M., and Labandeira, C. C. (2022). Insect herbivory on Catula gettyi gen. et sp. nov. (Lauraceae) from the Kaiparowits Formation (Late Cretaceous, Utah, USA). PLOS ONE, 17(1):e0261397.

Morris, R. J., Gripenberg, S., Lewis, O. T., and Roslin, T. (2014). Antagonistic interaction networks are structured independently of latitude and host guild. Ecology Letters, 17(3):340–349.

Novotny, V., Miller, S. E., Hrcek, J., Baje, L., Basset, Y., Lewis, O. T., Stewart, A. J. A., and Weiblen, G. D. (2012). Insects on plants: Explaining the paradox of low diversity within specialist herbivore guilds. The American Naturalist, 179(3):351–362.

Perez, T. M., Rodriguez, J., and Mason Heberling, J. (2020). Herbarium-based measurements reliably estimate three functional traits. American Journal of Botany, 107(10):1457–1464.

Rivera-Hutinel, A., Bustamante, R. O., Marín, V. H., and Medel, R. (2012). Effects of sampling completeness on the structure of plant–pollinator networks. Ecology, 93(7):1593–1603.

Schachat, S. R., Payne, J. L., and Boyce, C. K. (2023). Linking host plants to damage types in the fossil record of insect herbivory. Paleobiology. Early view.

Schachat, S. R., Payne, J. L., Boyce, C. K., and Labandeira, C. C. (2022). Generating and testing hypotheses about the fossil record of insect herbivory with a theoretical ecospace. Review of Palaeobotany and Palynology, 297:104564.

Schellenberger Costa, D., Zotz, G., Hemp, A., and Kleyer, M. (2018). Trait patterns of epiphytes compared to other plant life-forms along a tropical elevation gradient. Functional Ecology, 32(8):2073–2084.

Schleuning, M., Fründ, J., Schweiger, O., Welk, E., Albrecht, J., Albrecht, M., Beil, M., Benadi, G., Blüthgen, N., Bruelheide, H., Böhning-Gaese, K., Dehling, D. M., Dormann, C. F., Exeler, N., Farwig, N., Harpke, A., Hickler, T., Kratochwil, A., Kuhlmann, M., Kühn, I., Michez, D., Mudri-Stojnić, S., Plein, M., Rasmont, P., Schwabe, A., Settele, J., Vujić, A., Weiner, C. N., Wiemers, M., and Hof, C. (2016). Ecological networks are more sensitive to plant than to animal extinction under climate change. Nature Communications, 7(1):13965.

Swain, A., Azevedo-Schmidt, L. E., Maccracken, S. A., Currano, E. D., Dunne, J. A., Labandeira, C. C., and Fagan, W. F. (2023). Sampling bias and the robustness of ecological metrics for plant–damage-type association networks. Ecology. Early view.

Swain, A., Maccracken, S. A., Fagan, W. F., and Labandeira, C. C. (2022). Understanding the ecology of host plant–insect herbivore interactions in the fossil record through bipartite networks—Corrigendum. Paleobiology, 48:353–355.

Wappler, T., Labandeira, C. C., Rust, J., Frankenhäuser, H., and Wilde, V. (2012). Testing for the effects and consequences of mid Paleogene climate change on insect herbivory. PLoS ONE, 7(7).

Webber, Q. M., Schneider, D. C., and Vander Wal, E. (2020). Is less more? A commentary on the practice of ‘metric hacking’ in animal social network analysis. Animal Behaviour, 168:109–120.

Wilf, P. and Labandeira, C. C. (1999). Response of plant-insect associations to Paleocene-Eocene warming. Science, 284(5423):2153–2156.

Xiao, L., Labandeira, C. C., Dilcher, D. L., and Ren, D. (2022a). Arthropod and fungal herbivory at the dawn of angiosperm diversification: The Rose Creek plant assemblage of Nebraska, USA. Cretaceous Research, 131:105088.

Xiao, L., Labandeira, C. C., Dilcher, D. L., and Ren, D. (2022b). Data, metrics, and methods for arthropod and fungal herbivory at the dawn of angiosperm diversification: The Rose Creek plant assemblage of Nebraska, U.S.A. Data in Brief, page 108170.

